# Geometric Characterization of Pediatric Multiple Sclerosis Lesion Morphology: A Cross-Sectional and Temporal Analysis using Differential Geometry

**DOI:** 10.1101/2025.08.09.669512

**Authors:** Prateek Mittal, Prabal Pratap Singh, Joohi Chauhan

## Abstract

Pediatric-onset multiple sclerosis (POMS) involves aggressive inflammation and large lesions; however, its detailed shapes and structures remain poorly understood. Traditional volumetric metrics often overlook the complex geometry of these lesions. The main objective of this study is to develop a differential geometry-based framework to quantify the shape and structure of lesions in pediatric multiple sclerosis(MS) patients using longitudinal 3D FLAIR MRI. The goal was to identify reproducible lesion morphotypes and to track their evolution over time. Our approach sensitively tracked shape changes over time and revealed consistent progression patterns. Our findings suggest that geometric biomarkers offer a powerful new lens for decoding MS heterogeneity and tracking disease activity in the pediatric population.

## Introduction

Pediatric-onset multiple sclerosis (POMS), a form of MS affecting children and adolescents, occurs before the age of 18 years and accounts for nearly 3-5% of all MS cases (1). POMS is characterized by an aggressive disease course, higher relapse rates, and the presence of unusually large and diffuse brain lesions (2–4). Despite these severe clinical features, the detailed geometry of MS lesions in pediatric patients remains unclear. Existing MRI-based assessments rely overwhelmingly on lesion volume or count metrics that are blind to the complex morphological patterns underlying disease progression and treatment response (5).

Magnetic resonance imaging (MRI), particularly fluid-attenuated inversion recovery (FLAIR), is the imaging modality of choice for detecting MS lesions owing to its high sensitivity to white matter abnormalities (6). Conventional analyses reduce lesions to binary regions of interest, discarding crucial shape-related information that may reflect the underlying biological processes, such as axonal damage, remyelination, or inflammatory dynamics. In adult MS, early evidence suggests that lesion shape, curvature, and structural complexity may be correlated with pathology (7). However, these insights have rarely been extended to the pediatric MS population.

Emerging methods based on differential geometry and spectral shape analysis offer powerful tools to move beyond crude size-based descriptors. Metrics such as Gaussian curvature, shape index, and Laplace–Beltrami spectrum can encode the intrinsic properties of a lesion’s surface independent of orientation, position, or scale. When applied to lesion surfaces extracted from MRI, these features have the potential to reveal distinct morphometric signatures, capturing how lesions bend, evolve, and interact with their local tissue environments. Despite their promise, such geometry-based tools remain underutilized in studies of MS lesion morphology, particularly in pediatric cohorts.

In this study, we present a geometry-driven framework for analyzing lesion morphology in pediatric MS using 3D FLAIR MRI data. Using the publicly released PediMS dataset (8), which includes up to six longitudinal scans per patient, we extracted differential geometric and spectral descriptors from the lesion surfaces to construct a high-dimensional, rotation-invariant representation of each lesion. We then applied unsupervised clustering to identify natural groupings of lesions based on shape and tracked how individual lesions changed over time in terms of curvature, compactness, and spectral structure.

Our approach not only captures conventional morphometric variability, but also uncovers rare shape phenotypes, such as concentric ring-like lesions, which may indicate atypical or evolving subtypes of MS. By integrating cross-sectional and longitudinal geometries, we provide a richer lens for understanding pediatric MS heterogeneity, which is grounded in the physical structure of lesions, not just their size.

This study highlights the underexplored potential of surface-based lesion profiling in pediatric neuroimaging. We demonstrated that geometric features can stratify lesions into meaningful morphological subtypes and may serve as early indicators of lesion trajectory. Ultimately, our results suggest that shape-aware analysis could enhance clinical interpretation, improve monitoring strategies, and inform the development of personalized biomarkers for pediatric MS.

## Related Work

Conventional analyses of multiple sclerosis (MS) lesions have long relied on volumetric and intensity-based features. However, these metrics often fail to capture the complex morphologies that emerge in the disease process, especially in pediatric-onset MS (POMS), where the lesion shape and spatial dynamics may provide critical biological and prognostic information.

Recent advances in differential geometry have provided new tools for modeling anatomical structures in medical imaging. Surface-based features, such as Gaussian and mean curvature, compactness, and shape index, provide localized shape information that is robust to imaging orientation and scale. Spectral descriptors, particularly the Laplace–Beltrami spectrum, offer a compact yet global representation of surface geometry. Reuter et al. (9) first established this approach as a powerful tool for shape comparison, introducing the concept of “Shape-DNA” for biomedical surfaces.

The application of these tools in neuroimaging is increasing. Levenson et al. (10) demonstrated how curvature-based features augment traditional radiomics, while spectral signatures have been used in cardiac imaging for tracking deformation over time (11). However, their deployment in MS lesion analysis remains limited and virtually absent in pediatric cohorts. In adult MS, some efforts have moved beyond volume to include shape analyses. Newton et al. (12) examined over 1,000 lesions and found that MS lesions are significantly more asymmetric and morphologically complex than non-MS white matter lesions. Similarly, Oh et al. (13) linked lesion conformational changes, such as the loss of sphericity over time, to disease progression. However, these studies did not incorporate spectral or curvature-based descriptors and were rarely extended to temporal morphometry at the lesion level.

Unsupervised clustering approaches have also shown promise for stratifying MS pathology. Rúa et al. (14) used lesion-level MRI features and PCA-based clustering to identify clinically distinct subgroups of patients. However, their approach was limited to intensity and volume metrics without leveraging the shape geometry.

In the pediatric domain, Banwell et al. (15) and Peña et al. (16) characterized unique imaging patterns of POMS, such as higher lesion burden in the posterior fossa and greater gadolinium enhancement. However, to date, no study has systematically analyzed lesion shape in pediatric MS, despite its potential relevance in understanding neuroinflammatory dynamics in the developing brain.

Parallel to these efforts, the deep-learning revolution has accelerated lesion segmentation and classification. Reviews by Zeng et al. (17) and approaches by Cetin et al. (18) have achieved near-perfect segmentation metrics. However, these pipelines typically produce binary masks, offering little interpretability or insight into the shape-driven disease phenotypes.

Taken together, these threads underscore a significant gap:while tools from differential geometry and spectral analysis exist and MS lesion morphology is known to be clinically meaningful, no prior study has applied a comprehensive shape-based framework to pediatric MS lesions, especially in a longitudinal context. Our study bridges this gap by combining local and global geometric features with temporal tracking to uncover morphologically distinct lesion subtypes in POMS.

## Materials and Methods

### Dataset Description and Preprocessing

We conducted our study using the recently released **PediMS** dataset (8), a curated collection of 3D FLAIR MRI scans from nine pediatric MS patients (P1–P9), each with one to six longitudinal time points. In total, the dataset contained 31 scans capturing lesion evolution over periods ranging from weeks to years. Each volume *I*_*p,t*_ ∈ ℝ^*X×Y ×Z*^ is provided in NIfTI format along with a binary lesion segmentation mask *M*_*p,t*_, where *p* denotes the patient and *t* is the time point. Each lesion was assigned a unique integer label, which enabled instance-level tracking over time.

To ensure consistency across scans, all volumes were subjected to (i) histogram-based intensity normalization to a reference scan, (ii) skull stripping using FSL BET, (iii) isotropic Gaussian smoothing with a 1 mm kernel, and (iv) removal of lesions smaller than 20 voxels to exclude potential artifacts.

### Surface Mesh Reconstruction

For each lesion label within the consensus mask volumes, we extracted the corresponding binary segmentation and reconstructed a triangular mesh by using the `marching_cubes` algorithm from(19)with an isovalue threshold of *ℓ* = 0.5. To avoid boundary artifacts, lesion volumes were zero-padded by 2 voxels in all directions prior to meshing. The mesh consists of vertex coordinates and face connectivity arrays formatted for compatibility with VTK. No explicit post-processing was applied beyond the surface extraction, ensuring a consistent topology across the lesions.

### Geometric and Curvature Descriptors

The meshes were analyzed using `PyVista` to compute the following:

- **Surface area** (*A*) and **volume** (*V*_mesh_) from the polygonal representation,
- **Compactness**: 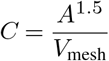.
- **Mean curvature** *H* and **Gaussian curvature** *K* per vertex using built-in curvature operators.

These curvatures are summarized using the mean and standard deviation across all mesh vertices.

### Spectral Shape Descriptors

The global lesion shape was encoded via the spectrum of the discrete Laplace-Beltrami operator, constructed as follows:

Triangle faces were iterated to compute cotangent weights (cot *α*, cot *β*, cot *γ*) for all edges,

- The cotangent Laplacian matrix *L*∈ ℝ^*n*×*n*^ and mass matrix *M* were assembled from vertex and face information,
- The generalized eigenproblem

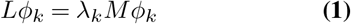

was solved using sparse Lanczos iterations with a shift-inversion strategy (*σ* = 10^*™*5^).

The first ten non-trivial eigenvalues *λ*_1_ through *λ*_10_ were retained per lesion. For normalization, eigenvalues were left unscaled, as compactness and volume were included separately in the feature set.

### Feature Extraction and Tabulation

Each lesion was summarized into a feature vector consisting of

- Surface area, volume (voxel- and mesh-based), and compactness,
- Mean and standard deviation of *H* and *K*,
- Laplace–Beltrami eigenvalues *λ*_1_ to *λ*_10_.

Only lesions with a minimum of 20 voxels were processed. For each lesion, we retained the patient ID, time point, and lesion ID for the longitudinal correspondence.

### Implementation Details

All computations were performed in Python 3.8 using:

- `nibabel` for NIfTI I/O,
- `scikit-image` for isosurface extraction,
- `PyVista` for curvature and mesh analysis,
- `SciPy` for Laplacian construction and eigendecomposition,
- `pandas` for feature aggregation.

Processing was parallelized per lesion and averaged for approximately 7 s per lesion on a 40-core Intel Xeon server with 256 GB RAM.

The full pipeline, including the mesh generation, feature extraction, and CSV output, is available in a reproducible opensource repository.

## Results and Discussion

### Unsupervised Shape Clustering

To investigate whether differential geometric descriptors capture meaningful lesion heterogeneity, we embedded lesion-level feature vectors into a 2D latent space using t-distributed stochastic neighbor embedding (t-SNE). The core hyperparameters for t-SNE were configured as follows: perplexity of 10, learning rate of 100, two output dimensions (n_components = 2), and a fixed random seed (random_state = 42) for reproducibility. The resulting manifold was clustered using HDBSCAN, where the hyperparameter min_cluster_size was set to five. HDBSCAN is a density-based algorithm that does not require a priori specification of the number of clusters. The embedding is visualized in Figure 1, where each point represents a lesion, and the colors denote the cluster assignment.

**Fig. 1.**
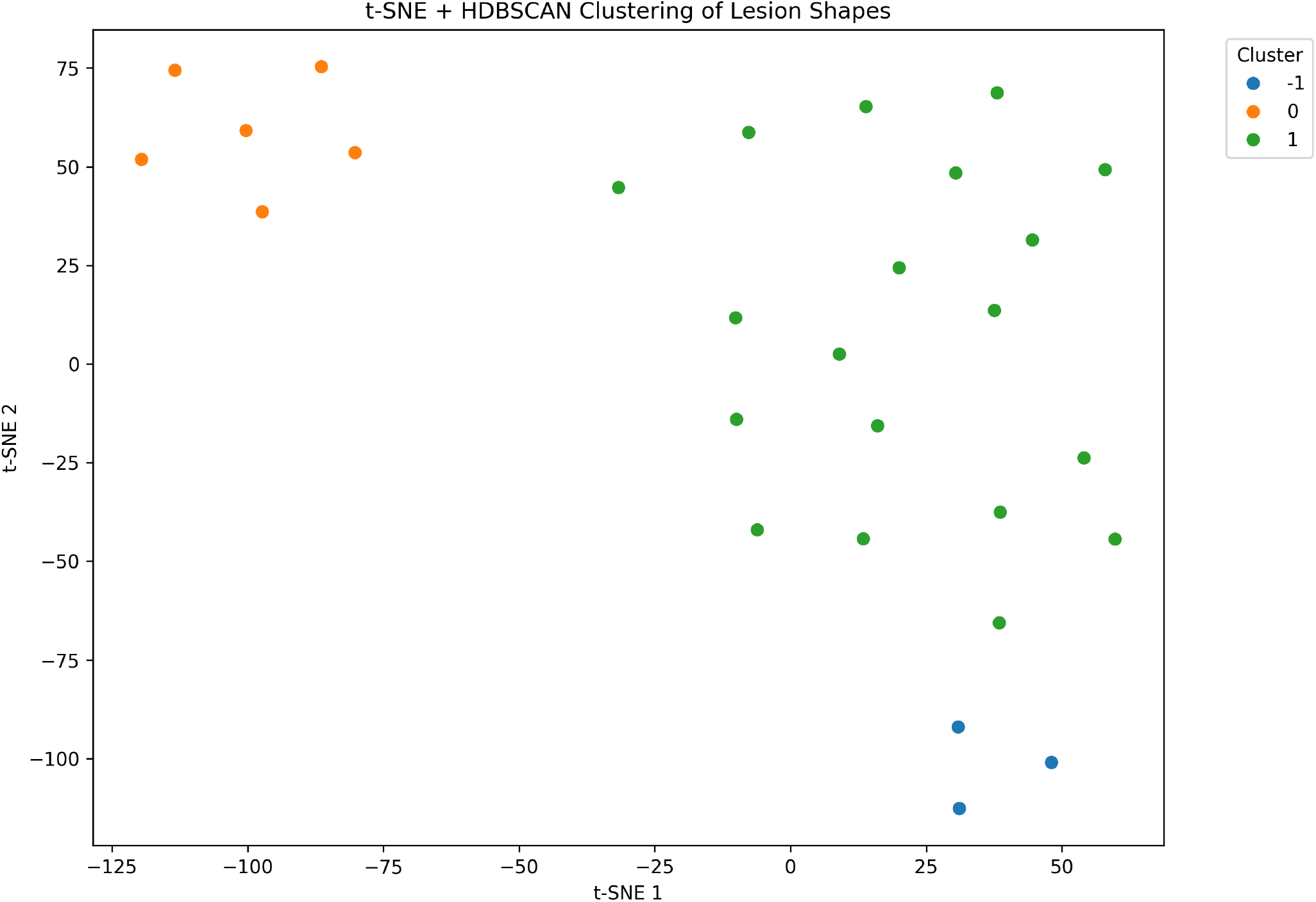
t-SNE visualization of lesion shape features clustered using HDBSCAN. Each point represents a lesion projected from 17-dimensional feature space. Colors indicate cluster membership. The clear separation suggests the presence of reproducible morphological subtypes.

Clustering quality metrics confirmed the presence of well-separated and internally consistent groups: a Silhouette Score of 0.523 indicated moderate to strong cohesion, a Calinski–Harabasz index of 28.18 reflected favorable inter-cluster separation, and a Davies–Bouldin index of 0.58 (lower is better), further supported by compact clustering. These findings suggest that shape descriptors encode latent, structure-driven subtypes that go beyond traditional lesion metrics such as volume or location.

### Morphological and Spectral Profiling of Lesion Clusters

To characterize the identified lesion clusters, we analyzed the distribution of the geometric and spectral descriptors (Figure 2).

**Fig. 2.**
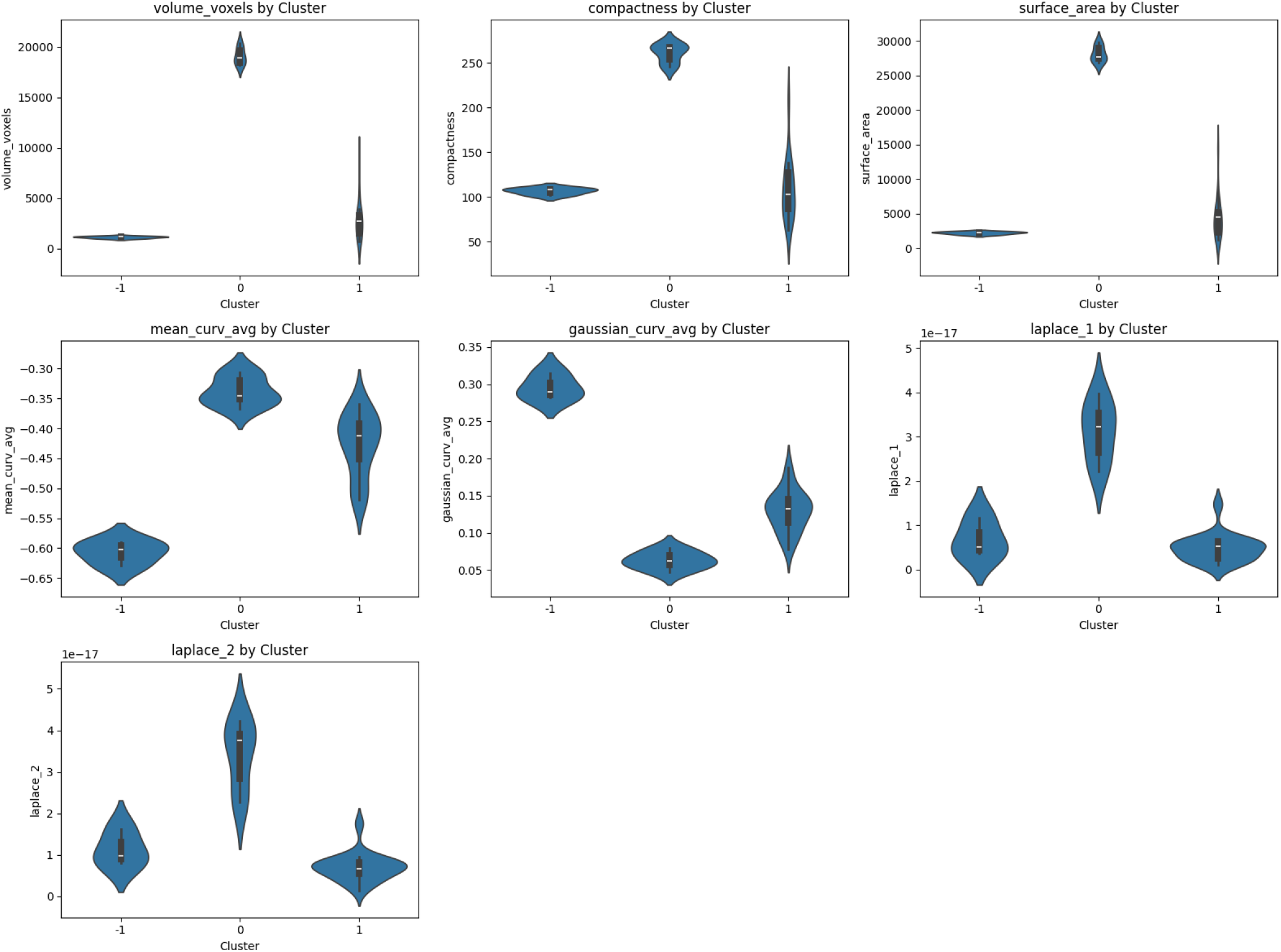
Violin plots showing the distribution of morphological features across lesion shape clusters. Key variables included lesion volume, surface area, compactness, mean and Gaussian curvature, and low-order Laplace–Beltrami eigenvalues. Distinct profiles highlight cluster-specific morphologies.

- **Cluster 0** lesions exhibited the largest volumes (mean: 19,091 voxels), high compactness, and smooth convex curvature, with relatively low spectral entropy. These properties may suggest structurally mature or possibly chronic plaques; however, confirmatory imaging (e.g., T1-hypointense mapping or contrast enhancement) is needed.
- **Cluster -1** comprised small lesions (mean: 1,115 voxels) with sharp curvature features — high positive Gaussian curvature and negative mean curvature — potentially reflecting newly emerging or resolving lesions. Their compactness and minimal surface area support this interpretation, although the temporal resolution remains a limitation.
- **Cluster 1** represented a heterogeneous group with high intra-cluster variability in shape, curvature, and spectrum. This cluster may span the intermediate or transitional lesion stages and capture a broad spectrum of lesions.

Differences across clusters were particularly evident in Laplace–Beltrami eigenvalue distributions, supporting the use of spectral descriptors complementary to traditional surface-based morphometrics. These spectral modes encode shape globally, and can stratify lesions into structurally and temporally distinct phenotypes without anatomical priors.

### Patient-Specific and Temporal Patterns

Next, we examined the distribution of lesion clusters across patients and at longitudinal time points. Cluster 1 lesions were found in nearly all patients and throughout the temporal axis, suggesting they represent common morphologies. Cluster 0 lesions were exclusive to patient P2 and increased in frequency from early to late scans (T1–T6), whereas Cluster -1 lesions were observed only in patient P7 during early stages (T1–T3). Figure 3 illustrates the temporal evolution of a representative Cluster -1 lesion, highlighting its spatial and structural stability across the three time points.

**Fig. 3.**
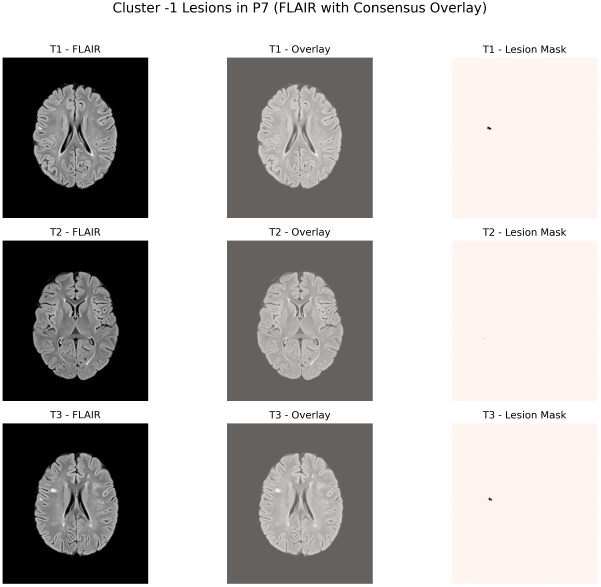
Longitudinal evolution of a Cluster -1 lesion in patient P7 across timepoints T1–T3. Each row shows the axial FLAIR slice (left), lesion overlaid on the FLAIR (middle), and binary mask (right). The lesion remains structurally and spatially stable with no appreciable growth.

The patient-specific distribution of these morphologies raises the possibility of underlying biological and clinical differences. Although clinical metadata were unavailable in this dataset, we hypothesized that Cluster -1 lesions may represent benign, non-progressive lesions, whereas Cluster 0 may capture chronic or coalescent plaques more commonly in progressive disease states. These hypotheses warrant further investigation using longitudinal clinical data, including EDSS scores, cognitive testing, and treatment status.

### Clinical and Developmental Implications

Pediatriconset MS presents unique neurodevelopmental challenges owing to the ongoing maturation of myelin, cortical pruning, and white matter tract consolidation. Lesion location and geometry during the critical windows of brain development may disproportionately influence long-term cognitive outcomes. Our finding that lesion-shaped clusters exhibit patient-specific and temporally consistent profiles suggests that geometric features could serve as individualized fingerprints of disease activity or recovery.

The persistence of compact, highly curved lesions (Cluster -1) may indicate early stage focal inflammation or resolution of demyelination, whereas the late emergence of high-compactness, low-curvature lesions (Cluster 0) in a single patient could represent cumulative pathology or treatment-refractory subtypes. These shape-derived signatures may offer greater interpretability and earlier indicators of disease progression than volume alone, particularly in pediatric cohorts, where volumetric comparisons are confounded by developmental variability.

## Conclusion and Future Work

This study introduces a differential geometry-based framework for the morphological profiling of multiple sclerosis lesions in pediatric patients using 3D FLAIR MRI. Unlike traditional volumetric approaches, our method captures intrinsic shape and curvature information through surface-based and spectral descriptors. Applied to the PediMS dataset, comprising nine patients and 31 longitudinal time points, our analysis revealed three distinct lesion clusters with robust geometric signatures and reproducible spatial and temporal patterns.

Crucially, we observed that lesion morphologies were not only distinct in shape metrics but also stratified by patient and time point, suggesting potential biological underpinnings. Cluster 0 lesions, which were larger, smoother, and possibly more chronic, were unique to a single patient and increased in prevalence over time. Cluster -1 lesions were compact, highly curved, and present only in one patient at the early stages. These findings suggest the possibility of shape-based phenotypes that can inform personalized monitoring and intervention strategies.

**Future work** should prioritize:

- Integration with clinical metadata (e.g., EDSS, cognitive scores, treatment history) to evaluate prognostic value.
- Validation in larger, multi-center pediatric MS cohorts to assess generalizability.
- Extension to other imaging modalities (e.g., contrastenhanced MRI, DTI, SWI) for multimodal lesion characterization.
- Investigation of lesion shape trajectories as early markers of treatment response or chronicity.

By establishing lesion geometry as a viable axis for disease characterization, this work lays the foundation for a clinically interpretable and developmentally informed paradigm in pediatric MS imaging.

